# SimpliPyTEM: An open-source Python library and app to simplify Transmission Electron Microscopy and *in situ*-TEM image analysis

**DOI:** 10.1101/2023.04.28.538777

**Authors:** Gabriel Ing, Andrew Stewart, Guiseppe Battaglia, Lorena Ruiz-Perez

## Abstract

Introducing SimpliPyTEM, a Python library and accompanying GUI that simplifies the post-acquisition evaluation of transmission electron microscopy (TEM) images, helping streamline the workflow. After an imaging session, a folder of image and/or video files, typically containing low contrast and large file size 32-bit images, can be quickly processed via SimpliPyTEM into high-quality, high-contrast .jpg images with suitably sized scale-bars. The app can also generate HTML or PDF files containing the processed images for easy viewing and sharing. Additionally, SimpliPyTEM has a specific focus on in situ TEM videos, an emerging field of EM, allowing for fast data processing into preview movies, averages, image series, or motion corrected averages using MotionCor2. The accompanying Python library offers many standard image processing methods, all simplified to a single command, plus a module to analyse particle morphology and population. This latter application is particularly useful for life sciences investigations. User-friendly tutorials and clear documentation are included to help guide users through the processing and analysis. We invite the EM community to contribute to and further develop this open-source package.

## Introduction

Electron microscopy (EM) is a powerful technique for observing samples at the nanoscale, and it is unrivalled for ease and popularity of use [1]. As with most modern-day microscopy methods, EM imaging nowadays yields data in the form of digital images or videos, or arrays of numbers, with high and low values representing bright and dark regions of the image respectively. In conventional, bright-field EM, the intensity of the signal corresponds to the density of the sample at each point, with more dense regions transmitting fewer electrons and thus yielding dark regions in the images. There is considerably more data available in the average EM image than can be seen with the naked eye, for example, the images are often 16-bit or 32-bit allowing for far more contrast than is displayed. Combined with this, there is often a considerable amount of noise in EM images, arising from various sources including from inelastically scattered electrons and electrons scattered due to multiple collision events [2], alongside undesired detector and readout signals. Noise can substantially obscure the underneath image. However, noise can often be reduced or removed using techniques ranging from simple linear filters [3] to complex methods, including deep learning-based methods [4-7]. Effective data processing of EM images is thus vital to maximising the information yielded by electron microscopy experiments. This requirement is even more significant in the world of *in situ* EM, where the ability to observe the behaviour of materials and nanoparticles under real-time, and controlled conditions can provide novel insights into advanced materials and novel nano-structures. *In situ* TEM typically generates a significant amount of data in the form of videos thus effective data processing and post-acquisition workflow optimisation become essential tasks.

With these requirements considered, a lot of specialised and manual image processing work is often required for post-experimental analysis. However, much of the post-experimental process cannot be easily automated, as the files produced are commonly incompatible with standard image viewing programs, they often have poor contrast and lack scale bars. As a result, many users spend significant time painstakingly performing basic post-imaging tasks including contrast enhancement, basic filtering and adding scale bars. These tasks can be automated. It can also be time-consuming to examine the acquired images as a whole or simply to produce a presentation to view and share these. Therefore, EM users are in constant need of methods to automate the basic data processing steps and allow the production of experimental contact sheets. Combined with this, the possibility to access the metadata from the images collected is also important, so users can quickly access details about images. However, metadata is often hidden within files and not easily accessible, making image curation a time-consuming task.

Many programs are available to process EM images, with one of the most common approaches being the use of ImageJ [8], an open-source package for scientific image analysis. ImageJ is effective for manually editing images and has a scripting-based macro language to automate repetitive tasks. However, this macro language has a steep learning curve, is poorly documented and not transferable to other systems. Gatan’s Digital Micrograph program [9] is also effective at various image analysis tasks, however is only available on Windows operating systems. Coded approaches can be very effective, the most popular language is Python. Many image analysis methods and modules are available with Python, including openCV [10], pillow [11] and scikit-image [12], making it possible to perform an enormous variety of tasks. One advantage of using Python for this analysis is the prevalence of Python-based machine learning and deep learning-based image analysis tools [13, 14], which commonly use images in a similar format. At the same time, the range of available options and locations can make it difficult for newcomers to locate the required methods. Many of the functions in these libraries also come with a large number of parameters which can make the function much more complicated to use than is necessary for most cases. The upside of using Python is huge however, Python is much faster for many tasks than ImageJ, and scripting can automate image analysis processes to reduce the amount of manual effort. While scripting tends to be more powerful than user-interface based approaches, many potential users find coding intimidating and challenging to learn, leading to users performing time-consuming manual methods.

Herein, we introduce a new app for basic image and video processing, and visualisation. In addition, we also introduce a Python library for work of added complexity. The app aims to allow effective image processing from large file-size images or video files in various formats, to reduced size, high-contrast JPEG files with scale bars. HTML or PDF documents containing these images and videos can also be created for easy visualisation. The Python library aims to build on available methods by lowering the barriers of users new to Python for analysing EM images and videos. An extensive range of functions, including image and video visualisation, filtering, contrast enhancement and scale-bar addition are available in a user-friendly, consistent and well-documented manner, with few required parameters. This package is accompanied by detailed documentation and iPython notebook-based tutorials to make it easy to access and follow for users with only basic knowledge of Python.

## Methods

### Image manipulation

The main aim of this package is to simplify the use of Python-based programming for processing and analysing electron microscopy images. Python already has many publicly available libraries for image analysis [10-12], some of which have been used to varying degrees in this package. Image processing with Python is commonly performed by holding images in a numpy array [15], these are n-dimensional matrices of values which store data and allow many efficient data-editing functions. These numpy arrays are the backbone of image handling and manipulation in SimpliPyTEM. OpenCV [10] is another key library used in this package, a popular image manipulation library containing many useful functions like image filtering with median and gaussian filters. These filters use 2D convolution to reduce the noise levels within the image and are commonly used when viewing noisy images. Other 2D image filters included that aim to reduce noise within the images, such as a Wiener filter, here implemented using SciPy [16].

SimpliPyTEM supports the opening and saving of various image and video format files. Digital micrograph (DM) [9] files, a common EM file format from Ametek (formally Gatan) detectors, are opened using openNCEM [17]. Metadata from DM files are extracted and easily accessible. Another popular EM image format, MRC files, are opened using the Python package mrcfile [18]. Both of these file types can include both images and movie files.

In-situ TEM experiments are often recorded with screen recording software due to insufficiencies in direct and charge-coupled detectors for capturing videos. However the data produced often require similar processing to detector-captured videos. Here, we support the opening of most major video formats, including MP4, MOV and AVI, these file types are opened using openCV’s VideoCapture module.

Several choices are available for outputting image and video files from the software. For images, there is a choice between TIF files and JPEG files. TIFs allow image data to be saved in the current conditions, thus producing uncompressed images or images with higher bitrates. The Python package tifffile is used for this task. JPEG files are also supported, in this case files are compressed, producing a much smaller file size. The generated images are high quality nonetheless and ideal for display purposes.

Movie files have many export options, allowing for various downstream applications. Movie frames can be saved as sequences of TIF Files or as a TIF format image stack. Image sequencing can be particularly useful when investigating a dynamic process via *in situ* EM. MP4 and AVI movie files are saved using moviePy; MP4 files are effective for viewing and displaying movies, in particular, the MP4 format was chosen for its suitability to be displayed on webpages. The AVI files produced are uncompressed or raw, this format was chosen to be compatible with ImageJ.

### Document generation

The project herein presented aimed to create an automated method to view, share and present data collected by electron microscopy. To achieve this aim, we opted to create PDF files containing all the images collected in an imaging session, this is a common format and easy to share as a standalone document. The Python package PyFPDF (https://pyfpdf.readthedocs.io/en/latest/) was used to generate the PDF documents. Furthermore, to allow the user a more interactive experience, an HTML file containing all the images and movie files from an experiment can be generated. An accompanying CSS stylesheet is also produced to add to the interactive viewing experience. The HTML is generated with the Python package Airium (https://pypi.org/project/airium/). These options generate documents which can act like photography contact sheets, allowing users to rapidly view images and identify the images suitable for further processing or presentation.

### Image and video plotting

The recommended usage method for SimpliPyTEM’s Python library is to use an iPython notebook, for example, a Jupyter notebook [19]. These notebooks allow an interactive coding interface where images and plots can be displayed, and the underlying code can be easily edited and rerun. This method also allows new code to be run independently of the code which came before it, with variables still saved. To display images and plots within iPython notebooks, Matplotlib is used, while MoviePy is used to display videos.

### Specifically written algorithms

The *clip_contrast* method is used to improve the contrast of an image or video by scaling the image to new white and black values. The advantage of this method, rather than other examples like openCV’s enhanced contrast or histogram equalisation, is that *clip_contrast* can provide reliable improvements without any user decisions, and thus can be used in automated pipelines. Image contrast is selected by entering maximum and minimum pixel values or a saturation value. The saturation here is the percentage of pixels above a maximum or below a minimum value. Hence the minimum and maximum values are selected using the saturation. The pixel values in the image are then scaled to the new minimum and maximum values, such that these values are between 0 and 255.

We have also implemented l*ocal_normalisation*, this aims to even the contrast out across an image, as often TEM images are bright in the centre and dark in the corners. This effect can be visually displeasing, but more significantly it can make image segmentation using thresholding challenging. This algorithm separates the image into n x n patches, and then scales these patches to the global median, such that each pixel in the patch is multiplied by the global-median / local-median. To reduce edge artefacts from the patches, padding can be used. Padding in this context is an overlap between adjacent patches, and the mean of shared pixels in overlapping patches is used in the final image. An example of this method being used can be found within the documentation (https://simplipytem.readthedocs.io/en/latest/Tutorials/MicrographAnalysisTutorial.html#Fixing-uneven-contrast).

### Addition of Scale-bar

The scale bar is chosen as a suitable size for the image by considering the size of the image. The colour is chosen to be either black or white based on whether the scale bar area has a significantly lower mean pixel value than the rest of the image. The pixel values in the specified position are changed using numpy. The scale-bar text is added using pillow [11], as this allows special characters, including ‘μ’, which is commonly used in scale-bars (for micron units: μm). The user can convert the scale between nanometers and microns with a single command, while other conversions can also be performed but do require a scaling factor or measurements to be included.

### Particle analysis

A basic particle analysis module is included within the package, this is designed to collect statistics on the particle morphology, including area, circularity, and maximum, mean and minimum diameters. The module includes methods to threshold particles, using tools available with openCV, and extract data from the particles into dictionaries or pandas databases. Such databases can then be used to plot figures in Python or export the data to CSV files.

Particles can be located by finding edges in the binary thresholded image, internal parts of the particle are filled in, and particles larger or smaller than set values are filtered out, particles on the edge of the image are also filtered out. Again these functions mainly use openCV and the Python package imutils. A labelled image can also be inputted to find morphology data, allowing users to locate particles using other available methods, for example object detection programs like StarDist [14]. Morphology data from particles are collected and returned as a Python dictionary. An additional option to measure each particle across many positions is also included. In order to perform this measurement, every pair of coordinates is considered, and when these coordinates make an angle of 180°±1° with the centre point, the measurement of diameter is counted. This procedure was used in an image of polymer nanoparticles and shown in **Figure 3** and can be found in more detail in a tutorial within the library’s documentation (https://simplipytem.readthedocs.io/en/latest/Tutorials/Particle_analysis_tutorial.html).

### Metadata

Image metadata can be useful but difficult to access with TEM files. As such, we have tried to make it more accessible by saving the image or video metadata from DM files into a comma separated value (CSV) file, showing many of the key values saved within the file. By doing so, a user could easily check which images were collected at a certain magnification, or when the images were acquired by looking at an automatically generated table. Unfortunately, at present, this task only works with DM files.

### Sample images

The sample images included in the manuscript were collected on a JEOL 2200 microscope with a Gatan K2 camera. These images include a range of stained polymer nanoparticles, amyloid fibres and gold nanoparticles.

## Results

### SimpliPyTEM-GUI

The GUI-based image processing app is designed as a simple tool for EM users to use during or post-experiment (**Figure 1**). Images and videos in a number of common formats, including digital micrograph and .TIF can be loaded, enhanced, and saved into .JPG images. Basic processing tasks can be performed on the images, including gaussian and median filters, the addition of a scale bar and contrast enhancement. Different options are also available for videos, including DM image stacks, .mp4 and .avi files to be saved as an average image, a video (.mp4 or .avi), a motion-corrected average (using motioncor2), a tif sequence (i.e. saved as individual tif files) or a tif stack. This app provides many options for *in situ* users to create effective previews of their videos.

**Figure 1:**
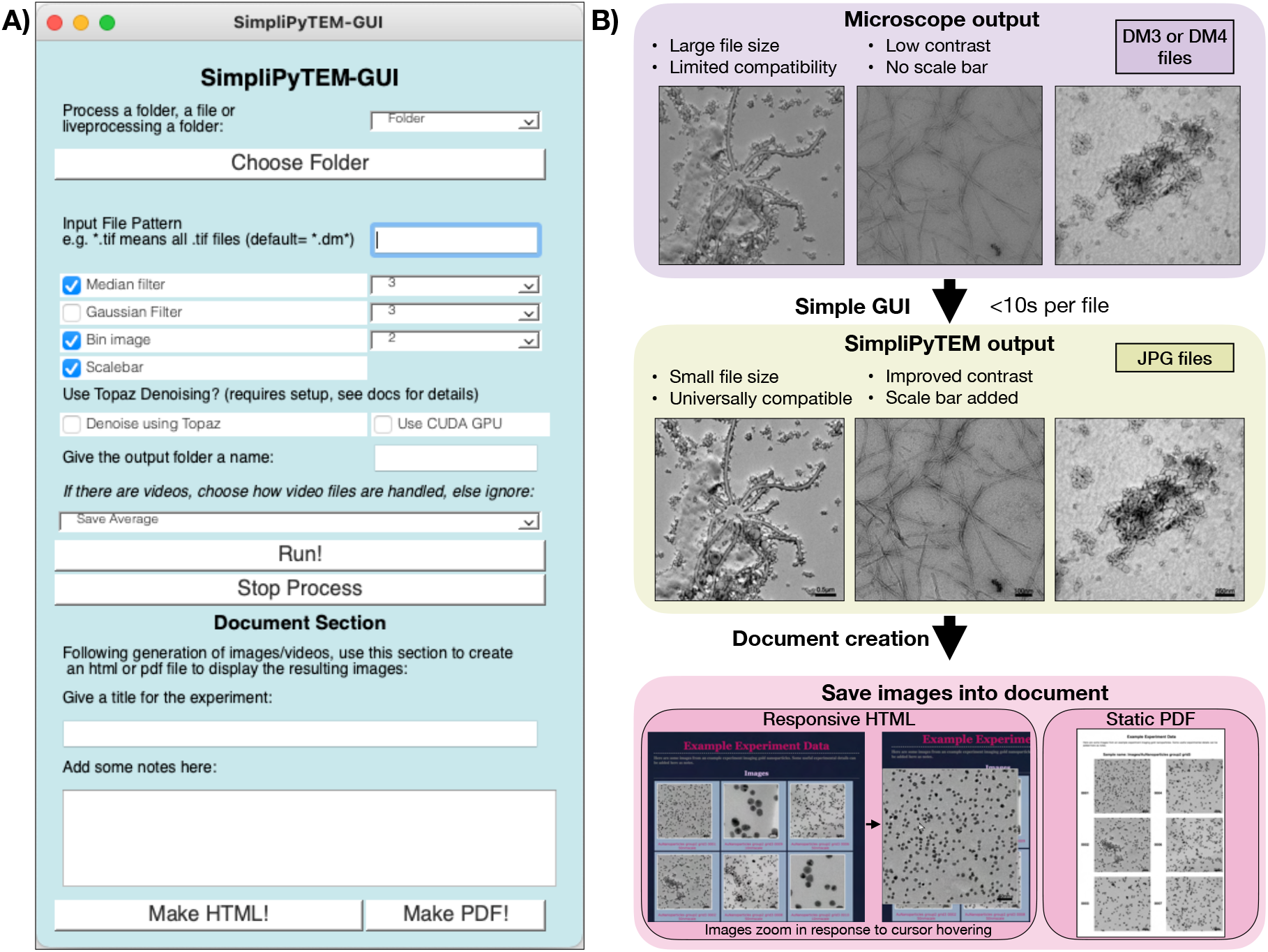
SimpliPyTEM-GUI for post-acquisition image processing. A) Appearance of SimpliPyTEM-GUI. B) Effect of SimpliPyTEM-GUI, showing the simple conversion of digital micrograph files with poor contrast and limited compatibility, to high contrast JPGs which can be used for observation and display. This process is quicker than many comparable methods, including using imageJ, taking seconds per file. There is also a document creation section which allows a PDF or responsive HTML document to be produced showing the images and videos collected during the experiment.

Image files from microscope detectors tend to be 16 or 32-bit and are unsuitable for regular uses, including adding them to a presentation or report. Due to the wide range of available pixel values present in the images, these also tend to have inadequate contrast as a default. This issue can lead to time being spent manually performing basic image analysis tasks. SimpliPyTEM-GUI can automate this process, by producing high-quality output jpeg files in only seconds.

In addition, SimpliPyTEM includes support for denoising images and videos using the deep-learning based denoising method Topaz [6]. Topaz can dramatically enhance images and videos, particularly by making low resolution features much easier to see. Although Topaz is specifically trained for cryo-EM, it can be highly effective for various bright-field images and videos. Unfortunately, this does have significant hardware requirements, needing a CUDA GPU for fast processing. By integrating Topaz within SimpliPyTEM-GUI, we aim to make state-of-the-art denoising methods more accessible and user-friendly for any researcher.

Alongside the image or video processing, there is also an additional section for visualisation. Images can be easily added to PDF or HTML files with a designated title and experimental notes. This process allows the rapid generation of documents to summarise the results of an experiment, which can be conveniently viewed or shared with others with very little preparation time.

As discussed in the introduction, image metadata can be useful for various reasons, for example to easily get an idea of the magnification used in an image. This value may provide information about what the acquired image contains without the need to view the image itself. Unfortunately, the metadata associated to images and videos’ is hidden within tags in DM files and thus can be difficult to access. However, SimpliPyTEM-GUI will automatically extract many of the features found within these metadata tags, including magnification, voltage, exposure time and acquisition date and time. These values are collected into a CSV table with the other files within the folder, which can then be easily examined, allowing easy identification of files by their imaging conditions.

### SimpliPyTEM-Library for images

Python is among the most powerful and widely used image processing and image analysis tools. There are lots of modules available for image analysis, which can be very effective, however many of these modules involve steep learning curves and unnecessarily large numbers of parameters. Therefore, to make common methods more available, we have created a library drawing upon some commonly used methods, including openCV, scikit-image and numpy [10, 12, 15]. This library is built on the principle of making the methods simple to use while sacrificing little performance. **Figure 2A** displays some of the available functions within the library. In contrast, **Figure 2B** shows a code snippet, demonstrating the simplicity of using the code, and **Figure 2C** shows the effect of the code snippet on a single example image. This code ran in less than 2 seconds on a MacBook Pro, 2018 for a 32-bit 3838×3710 pixel image, demonstrating the speed at which the processing can be performed.

**Figure 2:**
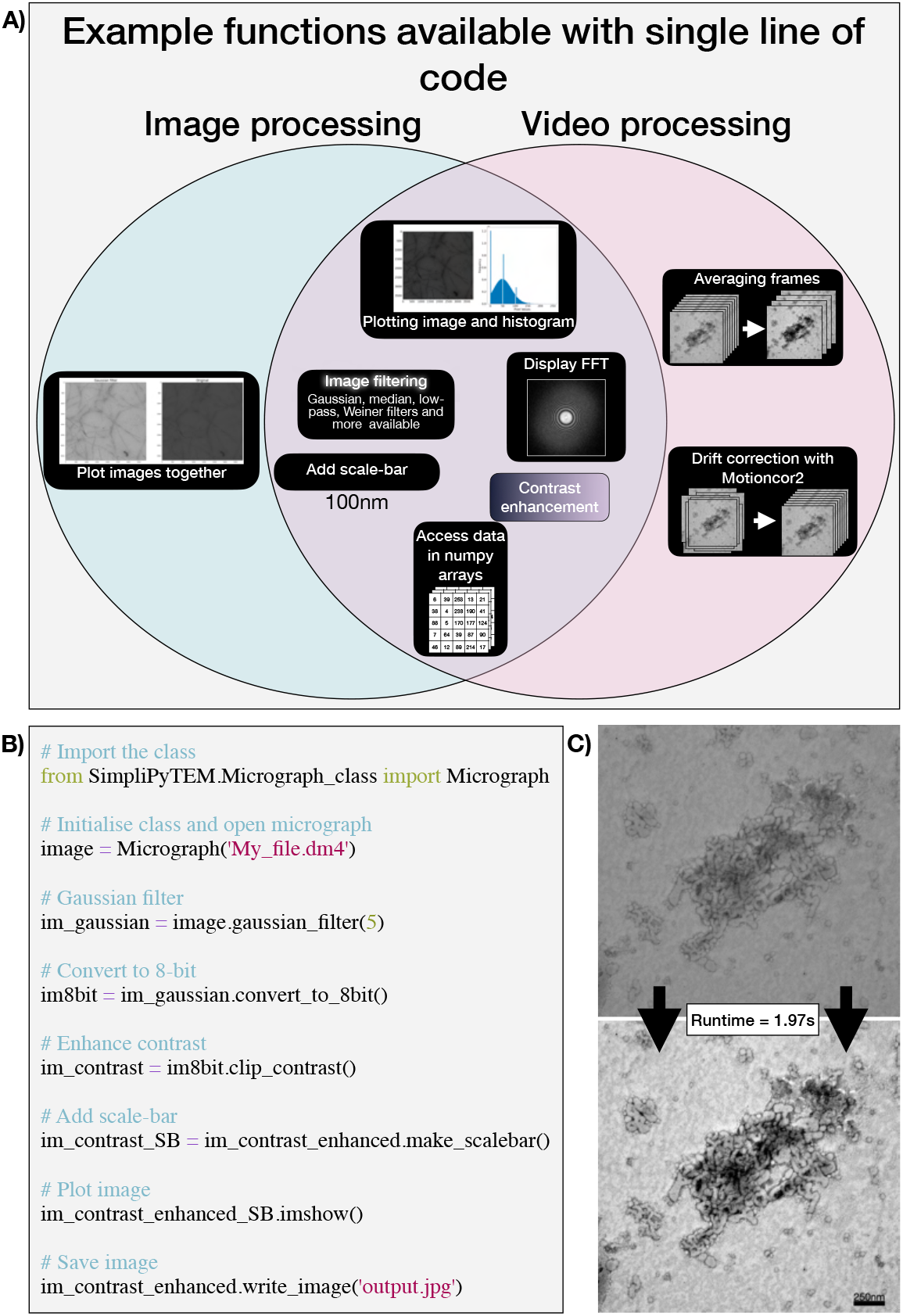
SimpliPyTEM - Python library. **A)** Mind-map showing an example of available functions that are all accessible with a single line of code. **B)** Example code for basic image processing. **C)** Image transformation achieved by the code shown in B, with a running time of 1.97s, with it falling to 0.84s when the image is not plotted (on MacBook Pro, 2018).

**Figure 3:**
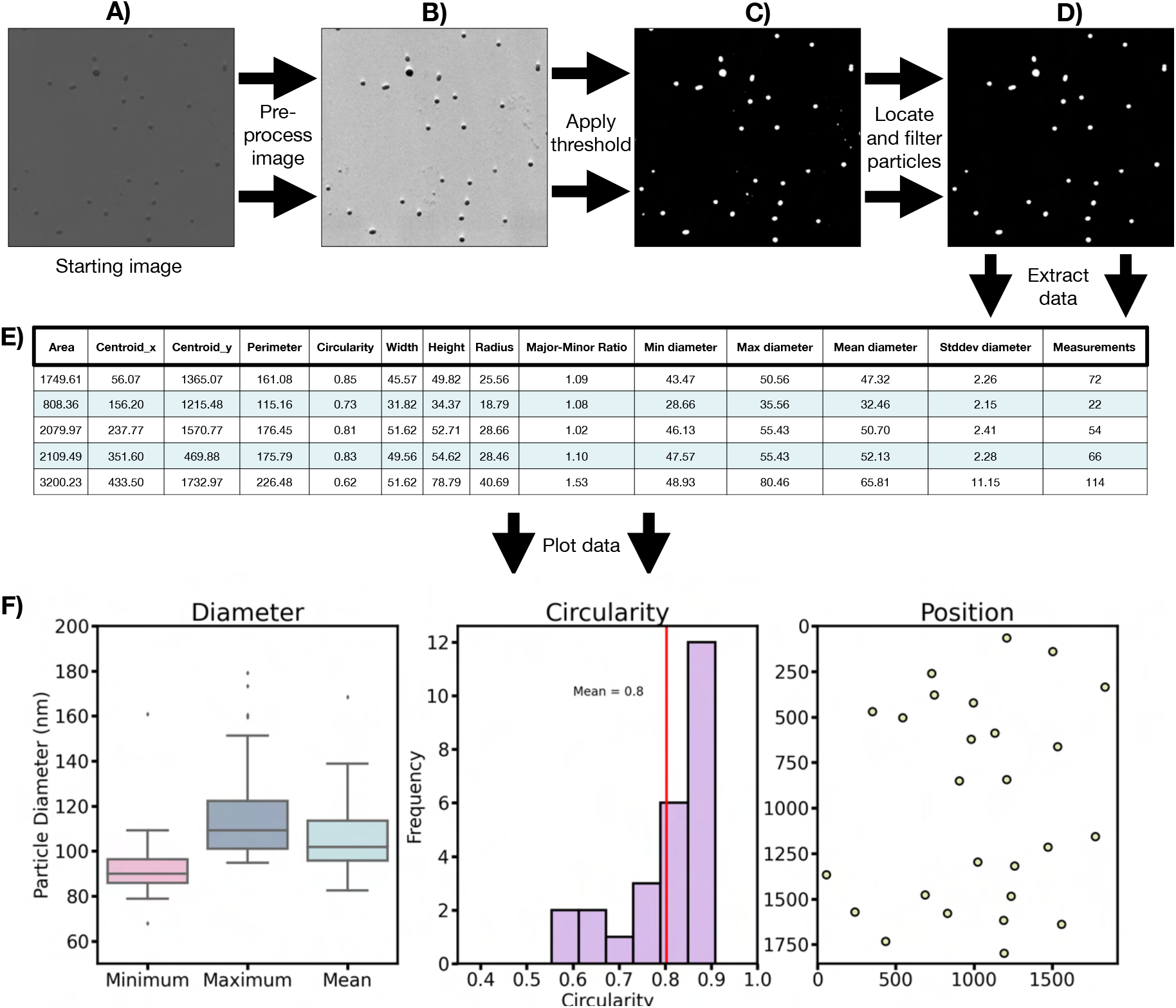
Basic Particle analysis protocol to extract particle data from a negatively stained image of polymer particles. The starting image is opened (A), preprocessed to enhance contrast and features (B), then a threshold is applied to create a binary image, with particles and background in white and black. respectively (C). The particles are located and filtered by size (D), particles touching the edges of the image are also removed. Finally, data is extracted (E) and plotted (F) with very few lines of code (from image to data). Here we show plots of particles’ diameter, circularity, and position, however more features are also accessible. The code to produce this analysis is available online as a tutorial within the documentation (https://simplipytem.readthedocs.io/en/latest/Tutorials/Particle_analysis_tutorial.html).

SimpliPyTEM image processing is primarily hosted in a single Python class called ‘Micrograph’. The Micrograph object hosts the image data, metadata and pixel size, and the methods to process the image. The library currently contains many simple methods to process the images, including image filtering (with median, gaussian, low-pass, non-local means and Weiner filters), the addition of scale-bar, converting to 8-bit, contrast enhancement and extracting metadata from digital micrograph images. In addition, images can be opened from various file types including digital micrograph (DM), MRC, TIF and JPEG files, and the edited images can be saved as TIF or JPEG files. These functions are all designed to be performed with a single line of code and with as few required parameters as possible, making the functions as easy as possible. These functions return a copy of the object, meaning the original object is kept.

### SimpliPyTEM-Library for videos

*In situ* TEM is a growing field, allowing the capture of live nanoscale events, providing dynamic information on phenomena that are not easily studied using other methods [20]. This results in EM videos containing a lot of information, particularly if captured from a direct electron detector, which commonly have high bit-rates and large sizes. Efficient analysis of acquired videos is essential, and as discussed in the introduction Python is ideal for this, however can prove difficult for inexperienced users. As such, we have created a Python library allowing for many basic and advanced methods to be employed for analysis of videos. Such methods include all of the techniques discussed for the image library, while also including many video-specific methods, for example, averaging frames together, video normalisation, and using existing software, motioncor2 [21] to correct for global motion within the video.

Videos can be loaded into a ‘MicroVideo’ object from various sources, including DM files, movie files like MP4, AVI, MOV and sequences of TIF or DM image files. From here, the videos can be averaged into groups of n frames, averaged in a sliding window fashion, converted to 8-bit, contrast enhanced and filtered, added a scale-bar, alongside several other functions. As with the image-processing library, these functions can be achieved in single lines of code, with few required parameters to make them as simple as possible. By processing videos in this way, the user can easily prepare the video for further analysis or presentations. Moreover, the video can be easily viewed in an iPython notebook (e.g., Jupyter notebook) and saved as an image sequence, an image stack, a single image (either single frame or average) or a movie file in .mp4 or .avi format, depending on its intended use.

### Particle analysis module

While the basic image and video processing modules, Micrograph_class and MicroVideo_class, are highly effective for image processing, we also include a basic image analysis module. The module contains simple methods for extracting data from nanoparticle, including positions, sizes, morphology, shape, and other physical properties. This information is crucial for thorough sample characterisation and optimisation in various fields, including pharmaceutical and materials science, nanotechnology, and biomedicine. The process for measuring particle properties is simple and involves applying a threshold to separate the nanoparticles from the background intensity, locating the boundaries of the particles and filtering the selected particles by area. This allows the user to measure the physical characteristics of the particles. These physical properties include area, position, circularity, major and minor axes, and major: minor axis ratio. The module also includes a way to take multiple measurements of the particle diameter from a single particle, allowing the diameter’s maximum, minimum, mean, and standard deviation to be collected. By considering these measurements, the user can obtain quantitative information about the uniformity of the investigated particles. An example of this functionality is given in **Figure 3**, where a negatively stained micrograph of polymer particles was selected for applying this module. Thresholding, was applied to the image, then the objects in the field of view, i.e. particles were located, filtered by size, and various properties could subsequently be extracted and plotted.

## Conclusion

Herein, we present a new computational Python package to aid with the processing and analysis of image and video data from electron microscopy experiments. The package is fully documented and supported by tutorials, aiming to make one of the most powerful image analysis tools more accessible to beginner users. The proposed package is particularly beneficial to *in situ* EM investigations where early data evaluation and post-processing can help users identify trends and correlations that may not be apparent from the raw data. In this fashion SimplyPyTEM allows users to make informed decisions about experiment design, sample preparation, etc by providing a fast and thorough evaluation of the data collected in the imaging session. Post-experimental image processing times can be reduced to mere seconds per file, and user-friendly documents to present, evaluate and share the data can be generated rapidly. By examining simple preview images within these documents, one can rapidly find the images or videos of particular interest for further analysis or display. Ultimately the aim for SimplyPyTEM is to share commonly used methods and unlock the potential of our data analysis. This, in turn, will help accelerate the science of all electron and *in situ* microscopy.

## Code availability

All the code is fully open-source and licensed under the GPL-3 licence. It can be downloaded and installed using Python’s package manager ‘pip’ (‘pip install SimpliPyTEM’), with the pyPI page for the package being found at https://pypi.org/project/SimpliPyTEM/. The code is also deposited on GitHub at https://github.com/gabriel-ing/SimpliPyTEM, documentation and tutorials can be found at https://simplipytem.readthedocs.io/en/latest/index.html.

## Acknowledgements

We would like to thank Valentino Barbieri, Chiara Cursi and Barbara Yus-Ibarzo, for providing samples which have become example images in this manuscript. GI acknowledges the Wellcome Trust for funding his studentship (222908/Z/21/Z).

